# Covalent conjugation of glucose oxidase on single-walled carbon nanotubes for glucose sensing

**DOI:** 10.1101/2024.10.14.618143

**Authors:** Hanxuan Wang, Subhasis Dehury, Melania Reggente, Vitalijs Zubkovs, Ardemis Boghossian

## Abstract

Glucose sensing and monitoring are crucial for biological and medical applications. Compared to existing methods, real-time detection and long-term monitoring are still required. Single-walled carbon nanotubes (SWCNTs) have excellent optical properties for sensing applications, which provide the possibility for designing a new generation of glucose sensors. In this study, we describe a method for covalently conjugate glucose oxidase (GOx) on SWCNTs as an optical glucose sensor. The functional groups are introduced by a photocatalytic reaction which acts as the handle for protein loading on SWCNTs. In this sp3 defect reaction, the optical properties of SWCNTs can be maintained. With a convenient bioconjugation reaction, the GOx could be covalently linked with SWCNTs. Compared to the non-covalent immobilization conjugates, the covalent conjugate sensor exhibits a much stronger optical response toward glucose, and the stability of the biosensor also increases in harsh conditions. At the same time, we also report the changing ratio of the original E_11_ and defected E_11_* peak during the bioconjugation reaction, which is also inspiring for reaction monitoring on SWCNTs.

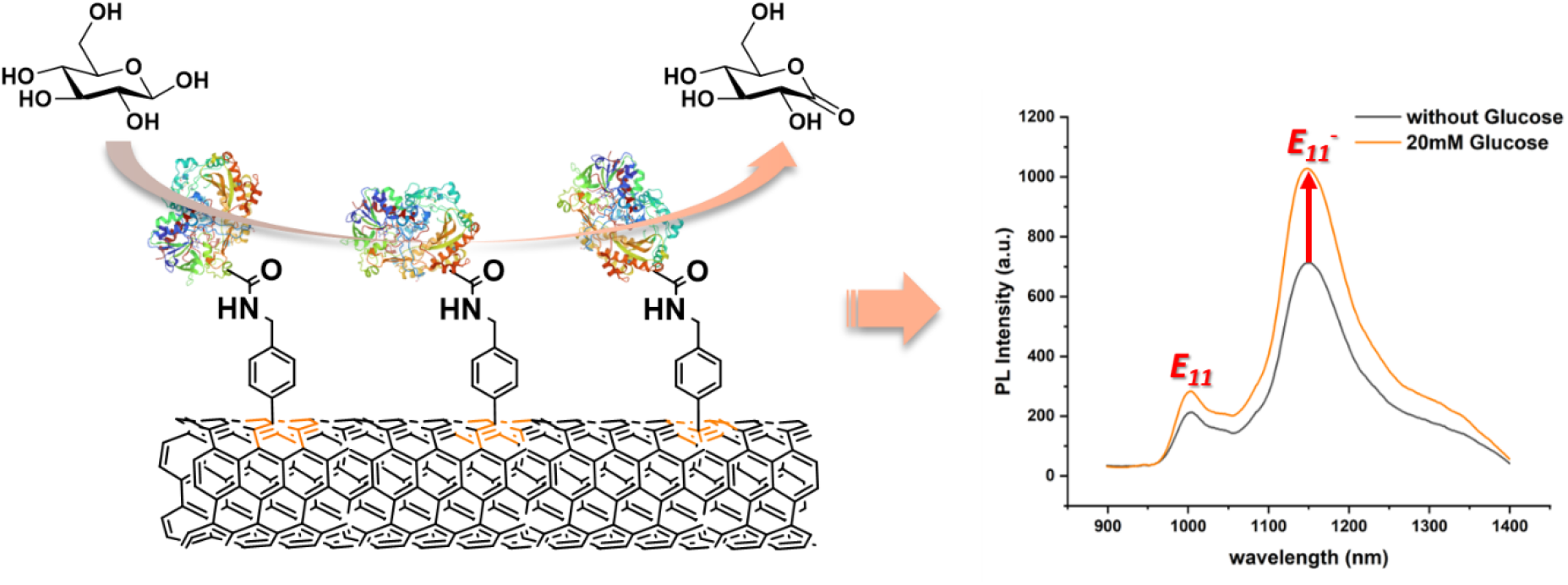

## 1. Introduction

Glucose plays a crucial role in the biological system, as it is considered to be the key molecule in cell metabolism.^1^ Glucose is involved in various pathophysiological processes and is fundamental to identifying certain health-related conditions.^2,3^ Testing and monitoring glucose levels is very useful when seeking to control diabetes, cell culture monitoring and food composition analyses.^4^ The progression of new technologies has seen measurement techniques of glucose levels become easier to use and more convenient.^5^ Most of these contemporary techniques use the electrochemical method, in which the sensors are expensive and not reusable.^6^ For example, the typical method to achieve the level of blood glucose is by fingerstick and then tested by a specific reader.^7, 8^ While this method can rapidly get a result, it can only be done at a single time point. For long-term and real-time monitoring, a new generation of glucose sensors is needed.^9, 10^ With excellent optical properties, SWCNTs show great potential in sensing applications.^11^ Single-walled carbon nanotubes (SWCNTs) may be conceptualized as a graphene sheet wrapped into a cylindrical form. Various protocols enable the preparation of SWCNTs with distinct chirality characteristics. Laser excitations see SWCNTs emit fluorescence that varies with the chirality and the environment of the SWCNTs, which are located at the second near-infrared (NIR-II) window.^12^ Emission wavelengths of the SWCNTs do not overlap with the maximum absorption wavelengths of hemoglobin, water, or other components distributed in blood and other biofluids and tissue.^13, 14^ Therefore, the fluorescence signal of the SWCNTs could penetrate biological samples with high optical stability. These advantages provide a genuine possibility to design new optical biosensors for glucose sensing.

To achieve a reliable fluorescence signal, a good suspension of SWCNTs is required. SWCNTs have a very low solubility in water, and surfactants such as sodium dodecyl sulfate (SDS), sodium cholate (SC) and sodium dodecyl benzene sulfonate (SDBS) are often used to suspend SWCNTs.^15-17^ SWCNTs could also be solubilized with biomolecules that could provide specific affinity toward analytes. DNA sequences,^18, 19^ RNA sequences,^20^ proteins,^21^ polysaccharides^22^ and lipids^23^ can be immobilized on SWCNTs’s surface for further sensing applications. For example, a collection of ssDNA sequences wrapped on SWCNTs could identify several target molecules such as neurotransmitters, tumor markers, and drug molecules.^18, 24, 25^ By monitoring the modulation of the NIR fluorescent spectra in response to the analytes, these ssDNA-SWCNTs could be applied as convenient biosensors for biological and pharmaceutical detections.

In addition to nucleotides, proteins or peptides could also be used to suspend SWCNTs.^21^ Research studies such as Vitalijs Zubkovs *et al*., developed a glucose oxidase (GOx)-based biosensor for glucose monitoring.^26^ Compared to DNA or RNA sequences, proteins have higher binding specificity toward the ligands, with ligands largely recognizing specific pockets of proteins. The interaction is based on Van der Waals forces, hydrophobic force, configuration changes, or energy transfers.^27, 28^ Therefore, the maintenance of the protein structures and activity is important for sensing applications. Previous work has focused on the non-covalent immobilization of enzymes on SWCNTs. Although the non-covalent conjugates are easy to prepare, these conjugates are heterogeneous. Non-covalent linkages are not as stable as covalent bonds, which might undergo degradation in physiological environments for in vivo applications.^29^ There have been several methods for covalent linking proteins on carbon materials, however, for sensing applications, the maintenance of the NIR signal must be considered.^30^ With sp3 defect chemistry on SWCNTs, the conjugated structure and original length of SWCNTs could be maximum retention.^31^ This is crucial for the NIR signal, with the functional groups potentially introduced at the same time. With these handles on SWCNTs, further bioconjugation could be achieved between proteins and SWCNTs.^32^ This new generation of SWCNTs chemistry provides the possibility to design covalent SWCNTs-based sensors.

Herein, we can report a photocatalyst reaction to link the iodine compound on SWCNTs, which provides the handle for protein crosslinking. During the course of this study, we were able to successfully link GOx onto SWCNTs. Compared to the non-covalent conjugate, the covalent conjugate shows a higher response toward glucose, and it demonstrated much better stability. We are also able to report changes in the ratio between the original E_11_ peak and E_11_* peak during the modification of SWCNTs. The results of this study suggest that this phenomenon could provide a new approach for real-time monitoring of complex reactions to SWCNTs.

## 2. Materials and methods

### 2.1 Materials

Purified SWCNTs were purchased from CHASM (SG65i, produced using CoMoCAT™ synthesis technology). Glucose oxidase (GOx) (from Aspergillus niger) was purchased from TCI. Bovine serum albumin (BSA) was purchased from Hello Bio™. 4-iodobenzylamine hydrochloride, 4-iodobenzoic acid, 1-Ethyl-3-(3-dimethylaminopropyl) carbodiimide (EDC), N-Hydroxysuccinimide (NHS), Tris(hydroxymethyl)aminomethane (Tris) and Sodium cholate hydrate were purchased from Sigma-Aldrich. Sodium Dodecyl Sulfate (SDS) was purchased from Carl Roth. Fluorescein isothiocyanate (FITC) was purchased from ABCR. PBS was purchased from Gibco™. Amicon® Ultra devices were purchased from Merck. Glass bottom Dishes were purchased from MatTek Life Sciences.

### 2.2 Preparation of SWCNTs suspension

45 mg of SWCNTs (SG65i) were mixed with 45 ml 1% SDS solution and homogenized for 30 min at 5,000 rpm (PT 1300D, Polytron). The mixture was sonicated in an ice bath for 1 hour (Q700 Sonicator, Qsonica, 10% amplitude). After sonication, the suspension was centrifuged at 10,000 RPM for 10 min at first and 30,000 RPM for 4 hours (Optima XPN-80, Beckman Coulter), and the supernatant was collected to remove the precipitates. To calculate the concentration of SWCNTs, we employed a UV-Vis-NIR spectrometer (Shimadzu 3600 Plus) to measure the absorption at 739 nm, and we calculated the concentration using an extinction coefficient of 25.3 mL/(mg·cm).

### 2.3 Sp3 defect chemistry on SWCNTs

The suspension of the SWCNTs was adjusted to a concentration of 40mg/L in 1% SDS. Deionized water was added to the suspension to further dilute the SWCNTs to 10mg/L. The 4-iodobenzylamine hydrochloride was dissolved in water and 4-iodobenzoic acid was dissolved in ethanol to 50mM, respectively. 50 µL of the solution was added to 5 ml diluted SWCNTs suspension with stirring. The reaction was excited by a 575 nm laser for around 48 hours (depending on the power of the laser and the amount of mixture) to form NH_2_-SWCNTs and COOH-SWCNTs. Green LED with higher power could be applied to larger-scale preparation. The fluorescence spectra of SWCNT were monitored during the reaction process. After the completion of the reaction, the mixture was transferred into the Amicon® Ultra device and washed with 1% SDS to remove the unreacted compounds. SDS could also be exchanged to SC via the Amicon® Ultra device to avoid the denaturing of proteins. The concentration of defect-containing SWCNTs was measured by UV-Vis-NIR spectrometer.

### 2.4 Loading of GOx on SWCNTs

The GOx protein was loaded on SWCNTs by covalent and non-covalent methods. For the covalent method, 20mg GOx was dissolved in water or the specified buffer. 1 μL EDC was dissolved in water, and 5mg NHS was dissolved in DMSO. If the sulfo-NHS (N-hydroxysulfosuccinimide) is applied, all the compounds could be dissolved in water. All the coupling reagents were added to the GOx solution for the activation. After 1 hour of mixing, the activated GOx was added to defect SWCNTs (NH_2_-SWCNTS) for the bioconjugation reaction. After the reaction, Amicon® Ultra devices were applied to remove the impurities to purify the conjugate. The non-covalent conjugates were using a previously-reported protocol (Zubkovs, V. et al, *Small* 2017, 13, 1701654). GOx was directly immobilized on both to original and defect SWCNTs through surface coating exchange in dialysis. These conjugates were used for further sensing and characterization experiments.

### 2.5 NIR fluorescence measurements

All the NIR spectra were obtained by a custom-built microscope setup (based on the Nikon Eclipse Ti-E microscope). Each well of 384-well plates was added with 20 μL of GOx-SWCNTs conjugates and 20 μL of glucose solution or deionized water was added. The SWCNTs were excited at 570 ± 10 nm (SuperK Extreme EXR-15 and SuperK Varia, NKT Photonics) laser, and the NIR fluorescence spectra were acquired between 950 nm and 1400 nm by InGaAs NIR camera (NIRvana 640 ST, Princeton Instruments) coupled with IsoPlane SCT-320 spectrometer (Princeton Instruments). Both the intensity of E_11_ and defected E_11_* peaks were measured to calculate the normalized response *I* towards glucose:

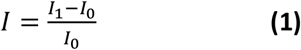

(*I* represent normalized intensity, *I*_*1*_ represents the intensity of E_11_ or E_11_* peaks of 20 μL glucose addition, with the final concentration of glucose 10mM, and *I*_*0*_ represents the intensity of the addition of 20 μL deionized water.)

### 2.6 Identify of colocalization of GOx and SWCNTs

The GOx protein was first labeled by FITC by the following procedure. Dissolved 10 mg GOx in 1mL NaHCO_3_/Na_2_CO_3_ buffer and 10 mg FITC in 100 μL DMSO. The solution of FITC was added to the GOx solution and the mixture was shaken at room temperature for 1h. After shaking, the mixture was stored at 4 °C overnight and purified by Amicon® Ultra devices. After the removal of free FITC, the absorption spectra of labeled GOx were obtained by Varioskan LUX Multimode Microplate Reader. The labeled GOx were linked with SWCNTs by the covalent or non-covalent methods described in **2.3**. All the conjugates were added to the poly-L-lysine-coated petri dishes and immobilized in the dark for 1 hour. After that, the petri dishes were washed with water and HEPES for three rounds (for the evaluation of stability, the petri dishes were washed with 5% SDS and methanol).

The co-localization of GOx and SWCNTs was identified on a custom-built microscope setup (based on the Nikon Eclipse Ti-E microscope) with oil-immersion TIRF Apo 100 x objective (N.A. 1.49, Nikon). The images were captured by InGaAs NIR camera (NIRvana 640 ST, Princeton Instruments) or EMCCD visible camera (iXon Ultra 888). All the image processing was done by Image Fiji software.

### 2.7 Zeta potential and FT-IR measurement

The suspension of SWCNTs was diluted in PBS buffer with a concentration of around 10 mg/L. Zeta potential was measured on Zetasizer Nano ZS (Malvern), with the DTS1070 cuvette.

### 2.8 FT-IR measurement

FT-IR measurements were performed in a Spectrum 3 FT-IR spectrometer (PerkinElmer). Samples were prepared prior to each measurement as follows. The suspensions of SWCNT-SC and SWCNT-NH_2_-GOx were drop casted on glass slides and heated for 10 minutes at 70 °C to evaporate water. Dry sodium cholate, GOx protein were directly put on the sample slide in a powder form. All spectra were recorded in triplicate for each sample types.

## 3. Results & discussion

### 3.1 Functionalization of SWCNTs

In literature are reported various methods for functionalization of carbon nanotube surface (REF). Most of these reactions are performed harsh conditions and includees treatments by strong acids, oxidation agents, or extremely high temperatures. These conditions might lead to the damaging of conjugation structure or fragmentation of SWCNTs which quench the fluorescence of SWCNTs. To reduce the damage and improve the conjugation were developed new generation sp3 defect chemistry. With a milder reaction system and better reaction control, the fluorescence of SWCNTs could be maintained for further applications.

As described in **2.3**, we employed a photocatalyst reaction with 4-iodobenzylamine hydrochloride to introduce the amino groups onto the SWCNTs (**Fig 1a**). Band gap changing led to the formation of the E_11_* peak around 1150nm.^33^ The intensity of E_11_* peak increases proportionally with the number sp^3^ defects on SWCNTs. The ratio between E_11_* and E_11_ peak could thus be used to index the reaction progression, the decrease of the E11 peak intensity was used to monitor the reaction progress and the reaction was considered to be completed when E11 peak intensity decreased by 80% from the initial intensity (**Fig 1b**). We also employed the PL (photoluminescence) map to evaluate the SWCNTs before and after the reaction (**Fig 1c**). From the PL-in map, we further confirm that the peak around 1150nm is originally from E_11_ peak of (6,5) SWCNTs rather than shift from other chiralities. In the acquired absorption spectra (**Fig S1**), the shape and position of peaks are similar, which provide the maintenance of the structure of SWCNTs.

**Figure 1.**
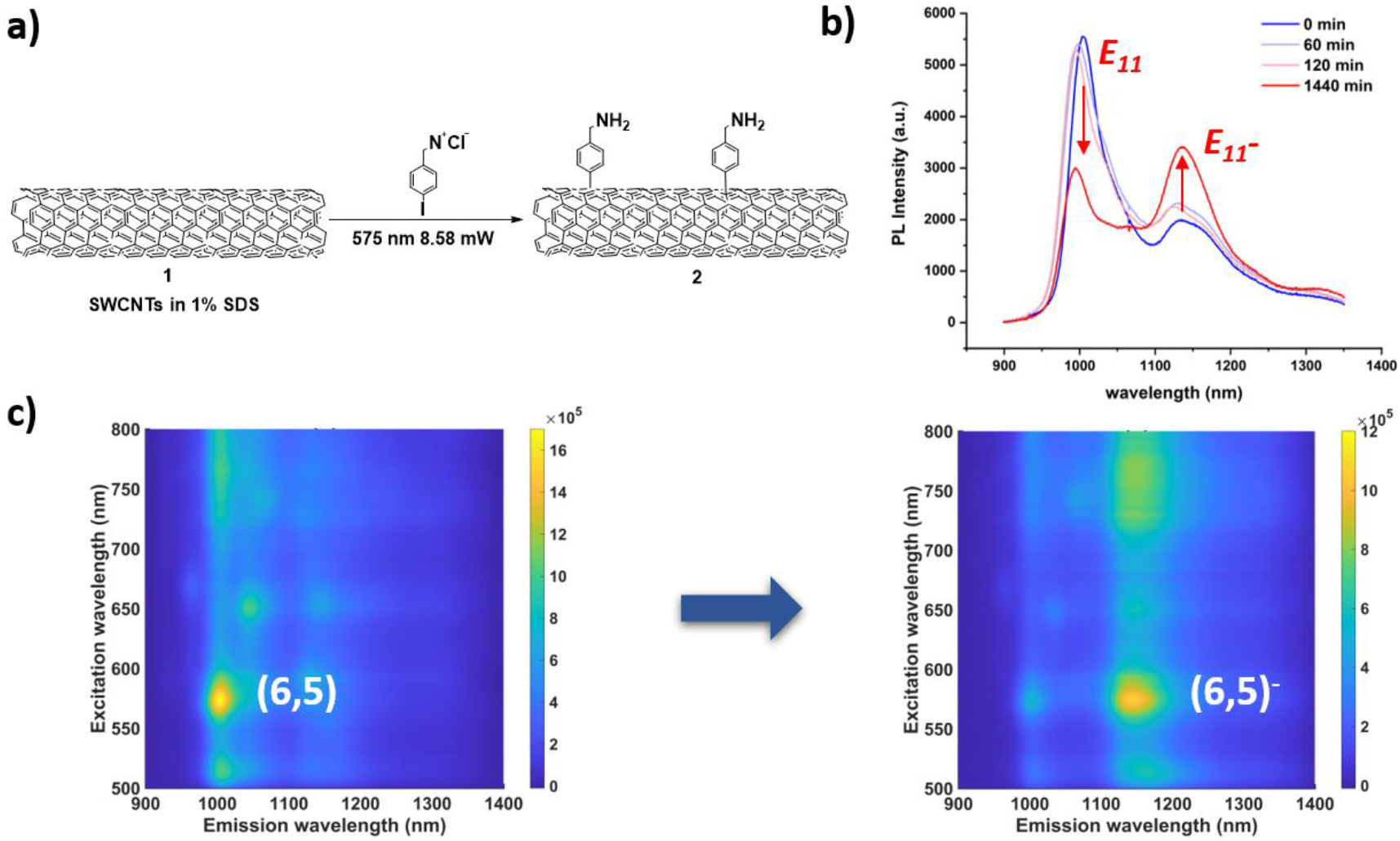
(a) Illustration of a reaction between SWCNTs and 4-iodobenzylamine hydrochloride. (b) NIR-fluorescence spectra of SWCNTs at different time points of the reaction. (c) PL map before (left) and after (right) the photocatalyst reaction.

After the successful demonstration of the sp3 defect reaction, we expanded the reaction to 4-iodobenzoic acid (**Fig S2**). Considering the amines are the most reactive amines for further condensation reaction, we prepared a larger batch of NH_2_-SWCNTs **2** for the next step of GOx loading. These reactions can be used to introduce amino groups and carboxylic groups under mild conditions and convenient purifications which is beneficial for bioconjugate chemistry on SWCNTs.

### 3.2 Loading of GOx on SWCNTs

With NH_2_-SWCNTs **2** in hand, we continued our attempts to covalently link GOx on SWCNTs. We screened different solvents of coupling reagents (**Fig 2a**), and we found with the existence of EDC, the reaction could happen. However, EDC itself has an interaction with SWCNTs, which leads to a strong increase in NIR intensity. This unusual rise covered other intensity changes and peak shifts and made the reaction difficult to monitor (**Fig S3**). Preactivated GOx with EDC and NHS provide the possibility to remove the excess EDC before adding to SWCNTs.^34^

**Figure 2.**
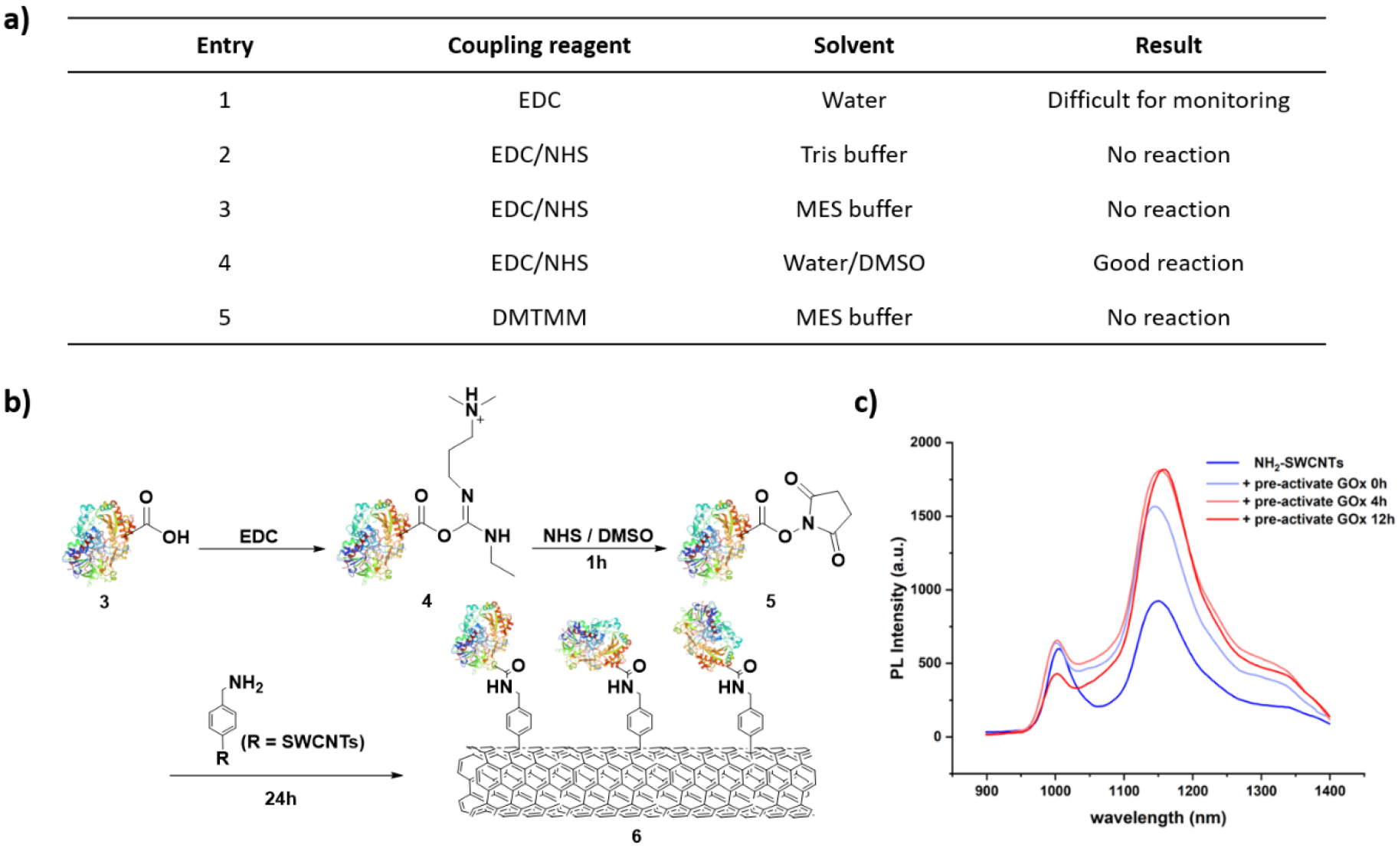
(a) Scheme of reaction conditions screening. (b) Illustration of reactions of GOx and NHS conjugation and the GOx linking with NH_2_-SWCNT. (c) NIR fluorescence spectra of NH_2_-SWCNT during its reaction with GOx-NHS.

We treated GOx with EDC and NHS in the DMSO/water cosolvent system and the formed active ester had moderate stability for ultrafiltration. After ultrafiltration, the excess coupling reagent can be removed, and the GOx with active ester terminal was mixed with NH_2_-SWCNTs (**Fig 2b**). We screened several buffers, we found that the ions in the buffer will affect the stability of SWCNTs and form precipitate. Therefore, the conjugation reaction still ran in the water. The reaction was then monitored by NIR spectra (**Fig 2c**). We found the E_11_ peak maintained the original intensity while the E_11_* peak raised a lot. The ratio between E_11_* and E_11_ changed from 1.54 to 4.24, this significant change showed the reaction site is in defect positions, which was also reported by Florian A. Mann *et al*.^35^ At the same time, redshift occurred at the E_11_* peak, which is also observed in many protein’s immobilization on SWCNTs protocols.^26^

### 3.3 Characterization of GOx-SWCNTs

After the synthesis of covalent and non-covalent conjugates, we further characterized the properties by FT-IR and Zeta potential. The vibrational mid-IR spectra if of SWCNTs themselves shows only few characteristic absorption peaks thus, monitoring the characteristic peaks of wrapping material on SWCNTs becomes important (**Fig 3a**). We noticed that the SWCNTs suspended in sodium cholate almost have the same spectra as pure sodium cholate powder. At the same time, we found in the covalent conjugate, a single peak around 1600 cm^-1^ disappeared and was replaced by the double peak of amide bound from protein.^36^ It is worth noting that, the shape of the double peak changed a lot, indicating that the conjugate is not a simple mixture, but with a stronger covalent link which affects the spectra of GOx. At the same time, the decrease of the C-H peak at 3000 cm^-1^ also shows the removal of sodium cholate.

**Figure 3.**
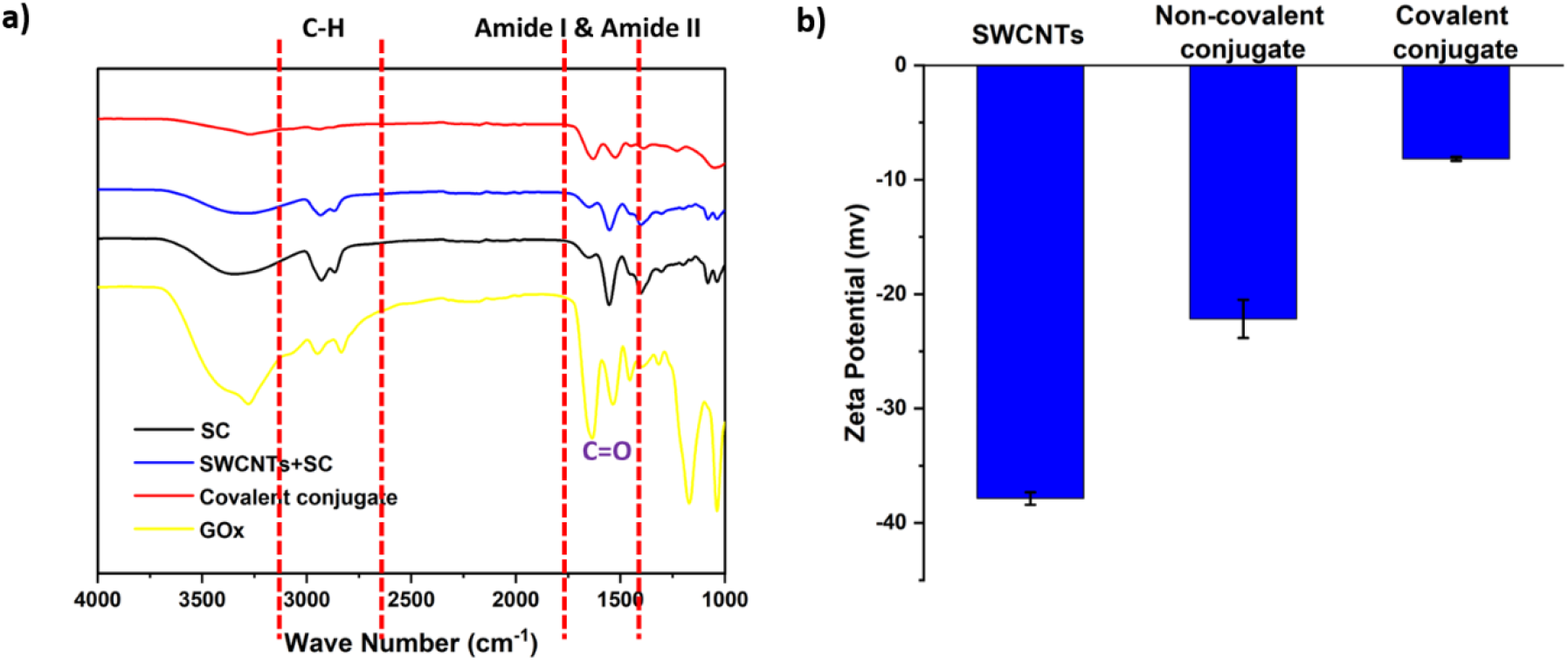
(a) FT-IR spectra of SC, SWCNTs in 2% SC, SWCNT-NH2-GOx, and XXX mg/mL GOx. (b) Measured zeta potentials of SWCNT-SC, SWCNT-GOx and SWCNT-NH2-GOx, respectively.

Measurements of a zeta potential was used to monitor changes in surface charge of SWCNTs’ (**Fig 3b**). The untreated SWCNTs suspension had a strong negative charge. With the immobilization of GOx, whether is non-covalent or covalent, the zeta potential increases a lot, this phenomenon is also reported with the immobilization of other proteins on carbon nanotubes.^37^ A higher zeta potential of GOx-NH2-SWCNT suggest higher protein concentration around surface of SWCNT compared to the non-covalent one. We believe that this result might be caused by the higher loading or better configuration of GOx on SWCNTs.

### 3.4 Colocalization of GOx and SWCNTs identification

To further identify the colocalization and evaluate the stability of the conjugates, we labeled GOx with FITC to visualize the proteins (**Fig S4**). With the same procedure with unmodified GOx, we prepared the covalent and non-covalent conjugates. Petri dishes were pre-coated with poly-L-lysine, and then we immobilized the conjugate on the surface of SWCNTs. After that, we washed the Petri dishes under very harsh conditions (5% SDS and methanol) to wash nonspecifically adsorbed GOx onto SWCNT surface, and then observed by microscope (**Fig 4a**). Both fluorescence of GOx and SWCNTs are stable under the treatment. The position of labeled GOx and SWCNTs still has a high consistency however, the non-covalent conjugate lost the colocalization between SWCNTs and GOx (**Fig 4b, 4c**).

**Figure 4.**
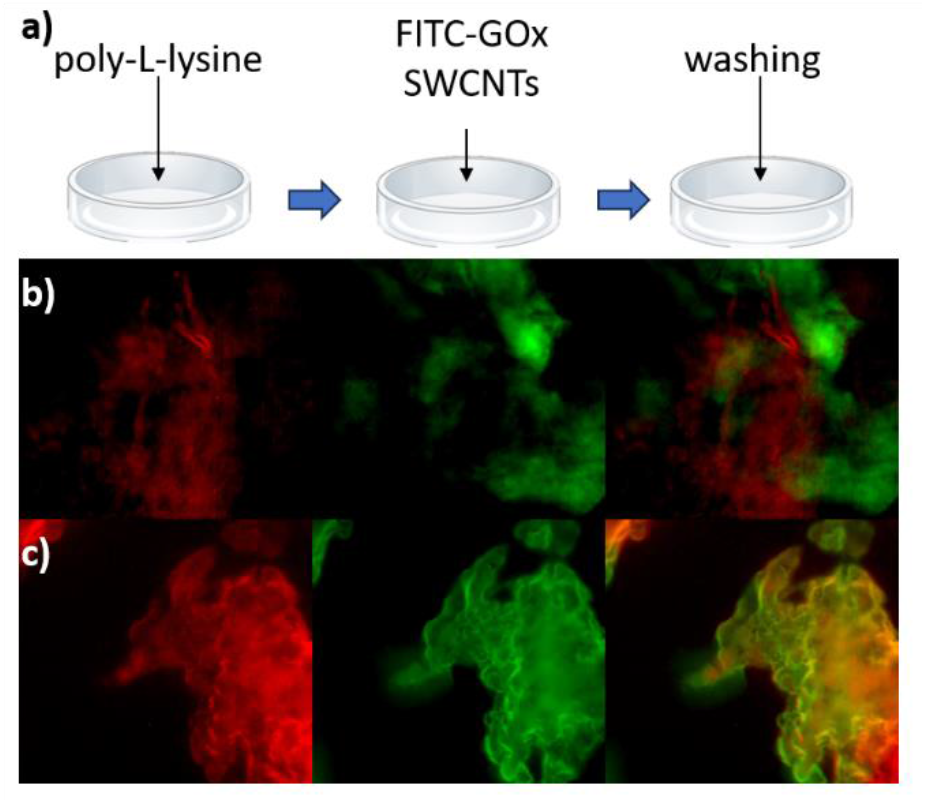
(a) Illustration of a sample preparation procedure before the microscopy experiments. (b) Noncovalent conjugate losses the colocalization of SWCNTs and GOx. (c) Covalent conjugate maintains the colocalization of SWCNTs and GOx.

The suspension of the covalent conjugate is also observed to have less precipitate than the non-covalent conjugate after adding methanol (**Fig S5**), which also proved the covalent bound can obviously increase the stability of the conjugate.

### 3.5 Evaluation of the response toward glucose

We have compared a sensory response for GOx-SWCNTs and GOx-NH2-SWCNTs by monitoring changes of SWCNT fluorescence intensity when interacted with 20 mM glucose solution in PBS (**Fig 5a**). A GOx-SWCNT sample was prepared according to a protocol previously published by our group. Where the sample was dialyzed against PBS for two days.

**Figure 5.**
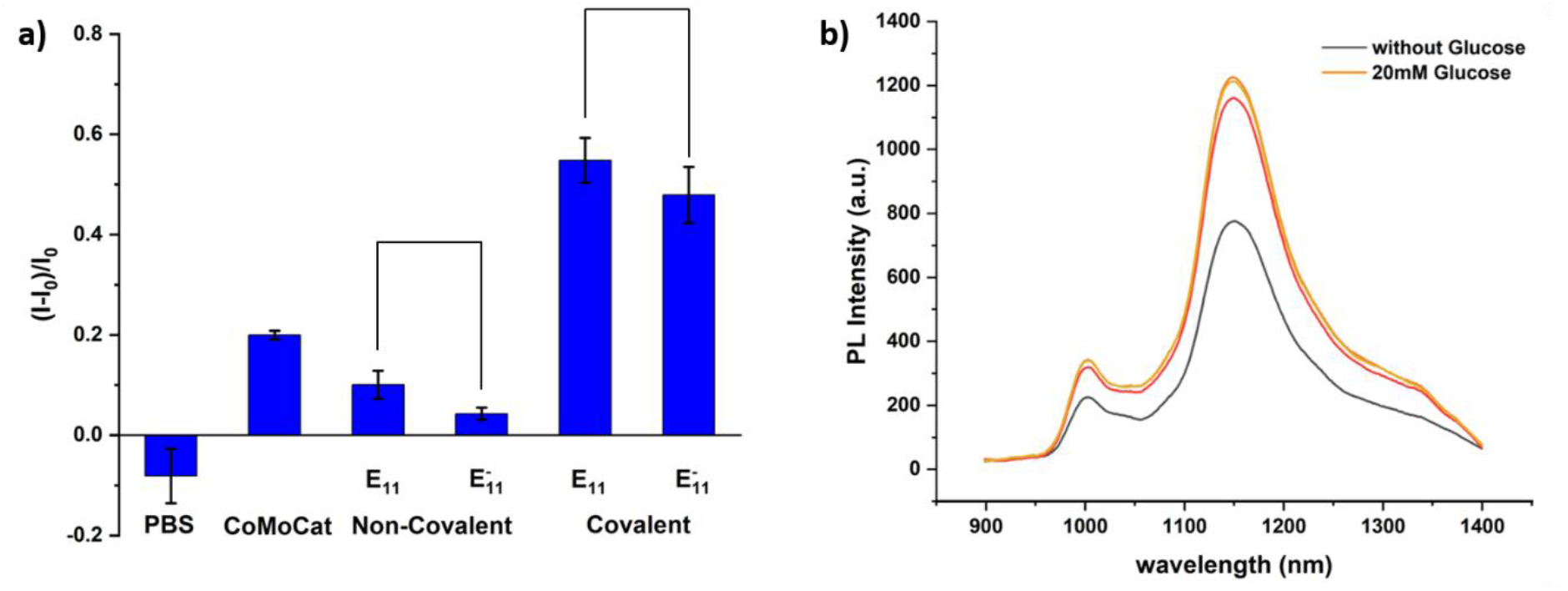
Optical response of SWCNT conjugates to glucose. (a) Difference of the fluorescence intensity before and XX minutes after addition of a glucose solution different SWCNT sensors. The E11 (6,5) SWCNT and E11* (6,5) SWCNT peak intensities were used for calculations (mean, n=3). The concentration of glucose is 20mM. (b) NIR-fluorescent spectra of GOx-NH2-SWCNT before and after adding glucose.

The GOx-(CoMoCat) SWCNTs showed an increase of fluorescence intensity by approximately 20% (Supplementary information equation **(1)**), We applied the same dialysis protocol for GOx immobilization on NH_2_-SWCNTs. For this case was observed that the increase of E_11_ and E_11_* peak peak intensities had a smaller magnitude compared to the GOx-SWCNT sample. We hypothesize that the functional groups affect the attachment of GOx on NH_2_-SWCNTs, and also makes it easier to form a precipitate as reported by Kruss.^38^ Surprisingly, the response on both E_11_ and E_11_* peaks of covalent conjugate become more significant and ***I*** can reach more than 50% intensity change (**Fig 5b**). A stronger signal from covalent conjugate suggests the higher GOx loading on SWCNTs. At the same time, we also notice that the E_11_ peak rises at the same level as the E_11_* peak, which might be because the covalent linking improved electron and energy transfer efficiency.

## 4. Conclusion

We have demotrated functionalization of SWCNTs with amino and carboxylic groups on the surface using sp3 defect chemistry. These functional groups behave as functional handles for further conjugation reactions. We demonstrated the application of GOx conjugation to NH3-SWCNTs. The resulted product showed greater sensitivity toward glucose. The covalent conjugate also shows better stability under harsh conditions, this also provides the possibility for further application in complex physiological environments. At the same time, we reported the changing ratio between E_11_ and E_11_* peak, this process can be real-time monitored by NIR spectra, which gives a new perspective on SWCNTs conjugation reaction monitoring.

## Acknowledgments

The authors are thankful for support from the Swiss National Science Foundation Project No. 200021_184822. This project has received funding from the European Research Council (ERC) under the European Union’s Horizon 2020 research and innovation program (grant agreement No 853005).

