## Supplemental information for "Covalent conjugation of glucose oxidase on single-walled carbon nanotubes for glucose sensing"

### 1. Absorption spectra

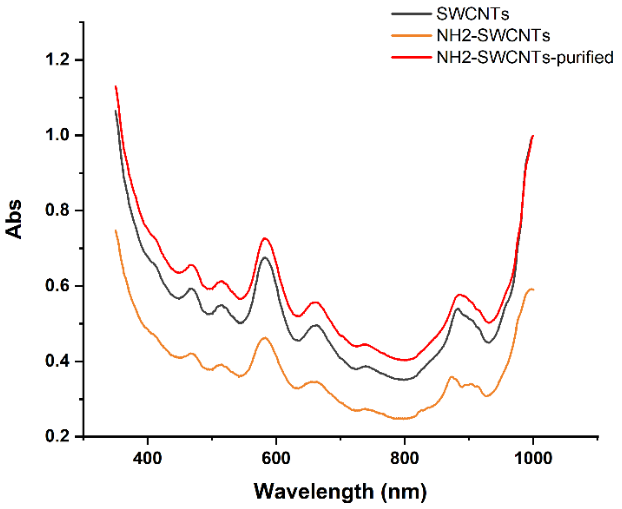


**Fig S1** UV-Vis-NIR absorption of SWCNTs before reaction (SWCNTs), after reaction ($NH_2$-SWCNTs), and after washing by water and 1%SDS to remove the impurities ($NH_2$-SWCNTs-purified).

### 2. Functionalization of SWCNTs

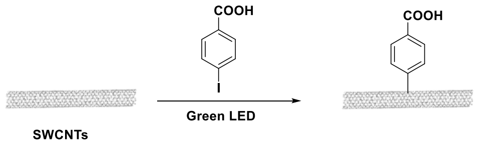


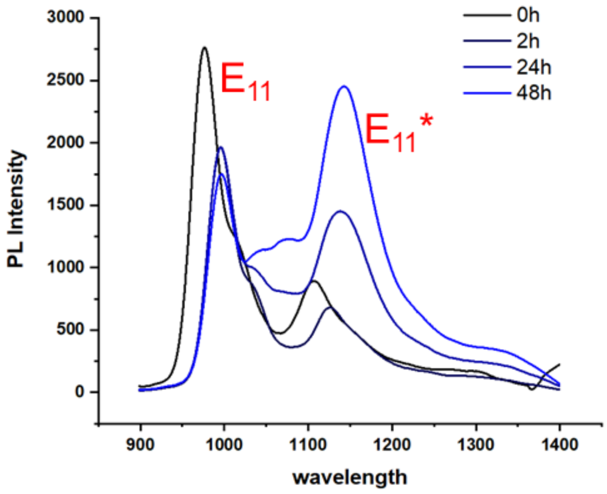

**Fig S2** Introduction of carboxylic groups on SWCNTs. Top: scheme of reactions, Bottom: Reaction monitoring by NIR-fluorescent spectra of SWCNTs.

### 3. EDC activation of $NH_2$-SWCNTs

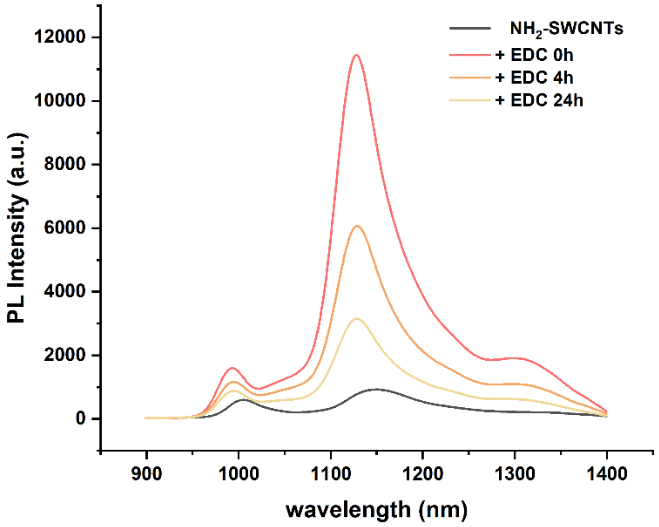


**Fig S3** NIR-fluorescent spectra of $NH_2$-SWCNTs. The $NH_2$-SWCNTs have a response toward EDC and cover the real shift from the reaction process.

### 4. Labelling of GOx

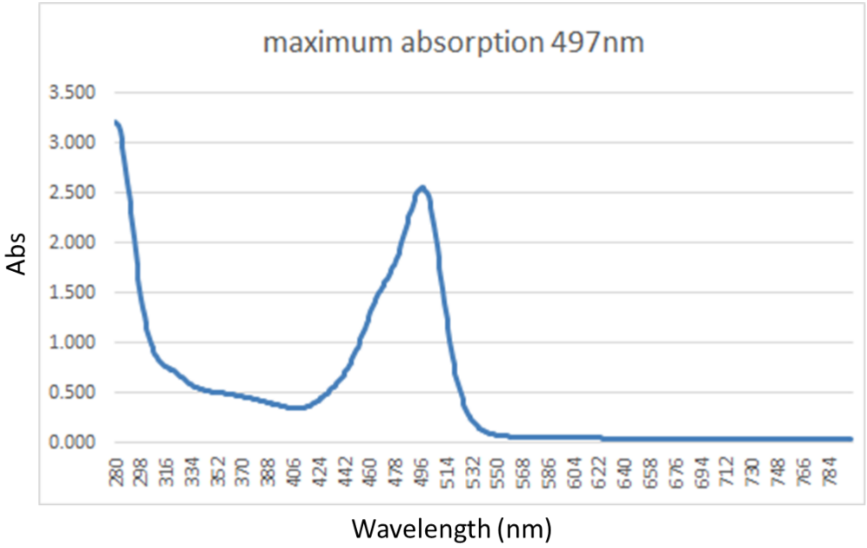


**Fig S4** Absorption spectrum of FITC-GOx, the characteristic peaks at 280nm and 497nm belong to GOx and FITC, respectively.

### 5. Stability of Conjugates

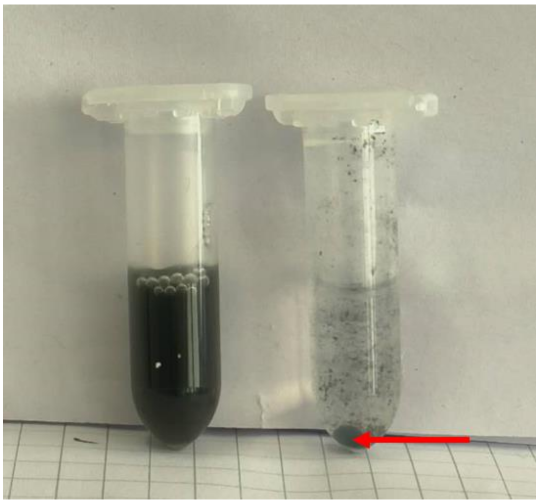

**Fig S5** Covalent conjugate and non-covalent conjugate after adding methanol. The covalent conjugate can still have suspension (left) and the non-covalent will form the precipitate very fast after centrifuge (right).
